# Mitochondrial accumulation and lysosomal dysfunction result in mitochondrial plaques in Alzheimer’s disease

**DOI:** 10.1101/2025.02.19.639081

**Authors:** Xiuli Dan, Deborah L. Croteau, Wenlong Liu, Xixia Chu, Paul D Robbins, Vilhelm A. Bohr

## Abstract

Dysfunctional mitophagy is a key component of Alzheimer’s disease (AD) pathology, yet direct *in vivo* evidence and mechanistic insights remain limited. Using a mitophagy reporter in an AD mouse model (*APP*/*PSEN1*/mt-Keima), we identified mitochondrial plaques (MPs) composed of accumulated mitochondria within or outside lysosomes in AD, but not normal mouse brains. Similar structures were also found in AD human brains, but not in healthy controls. Abnormal mitochondrial accumulation in dystrophic neurites, defective mitophagy, and impaired lysosomal function disrupted proper mitochondrial degradation, resulting in excessive mitochondria accumulation both within and outside autophagic vesicles. The resulting intensive mitochondria-containing neurites coalesce into MPs, which co-develop with amyloid plaques to form mixed plaques. These findings establish MPs as novel pathological entity and a promising therapeutic target in AD.

## Main text

Alzheimer’s disease (AD) is a devastating neurodegenerative disorder affecting millions of individuals worldwide, characterized by beta-amyloid (Aβ) plaques, neurofibrillary tangles (NFTs), and neuronal loss (*1*). The amyloid hypothesis suggests that Aβ accumulation drives AD pathology and cognitive decline, leading to FDA approval of anti-Aβ antibodies for early-stage AD. However, these therapies offer only limited benefits (*2*), and AD remains incurable (*1*). With the modest success of amyloid-based clinical trials, alternative mechanisms, including neuroinflammation (*3*), alterations of glial cells (*4*), cellular senescence (*5*), oxidative stress (*6, 7*), synaptic dysfunction (*8*), mitochondrial dysfunction/alterations (*9, 10*), and autophagy dysfunction (*11*) have been pursued as potential therapeutic targets. Recent findings consistently suggest the abnormal accumulation of autophagic vesicles and impaired lysosomal function in AD (*11–14*), but the specific substrates in these autophagic vesicles remain unidentified, limiting any potential therapeutic targeting. Early ultrastructural investigations showed a prominent presence of degenerating mitochondria in AD patients’ brains and our recent research (*15*) indicates impairment of mitophagy, a process that clears damaged mitochondria. Thus, we hypothesized that abnormal mitochondrial accumulation in AD may be linked to dysfunction of mitophagy, abnormal accumulation of autophagic vesicles, and amyloid pathology in AD. Due to the low spatial sensitivity and accuracy, previous indirect measurements of mitophagy impairment in AD mice (*15*) offer limited insight.

To facilitate direct measurement of mitophagy in specific brain regions, we established a mitophagy reporter transgenic AD mouse model, *APP*/*PSEN1*/mt-Keima, in which the pH-sensitive fluorescent reporter, Keima, is expressed in mitochondria. At the physiological neutral pH of the mitochondria, Keima has a shorter-wavelength excitation (green; 458 nm). Within the acidic lysosome, mt-Keima undergoes a gradual shift to longer-wavelength excitation (red; 561 nm) (*16*). As reported previously (*16–18*), the Keima excitation spectrums partially overlap. Consequently, merged images of the two channels show mitochondria in acidic environments as orange, while mitochondria in neutral compartments appear green.

### Mitochondrial plaques (MPs) are detected in mice and in human AD brains

Imaging fresh brain sections in the APP/PSEN1/mt-Keima mice at 50–60 wks showed that structures with abundant acidic mitochondria specifically formed in the cortex (Fig. 1A) and hippocampus of AD mice (Fig. S1A), but not in WT mt-Keima mice. Since these mitochondrial-enriched structures accumulate in a plaque-like format, we termed these structures mitochondrial plaques (MPs). The size of MPs ranged from several hundred to over a thousand square micrometers (Fig. 1B), with significantly smaller but more MPs detected in the cortex than hippocampus in AD mice (Fig. 1B, C). MPs exhibit distinct mitophagy signals compared to basal mitophagy, suggesting a pathological divergence from normal mitochondrial turnover. MPs are multilayered and contain bulbous-shaped subcomponents ranging from a few to over a hundred square micrometers (Fig. 1D-c) while basal mitophagy typically appears as evenly distributed spots under one square micrometers around cell nuclei (Fig. 1D-a, b). Approximately half of the MPs lack nuclei, while the remainder contain one or more (Fig. 1E, S1B), likely reflecting glial or neuronal cells.

**Fig 1.**
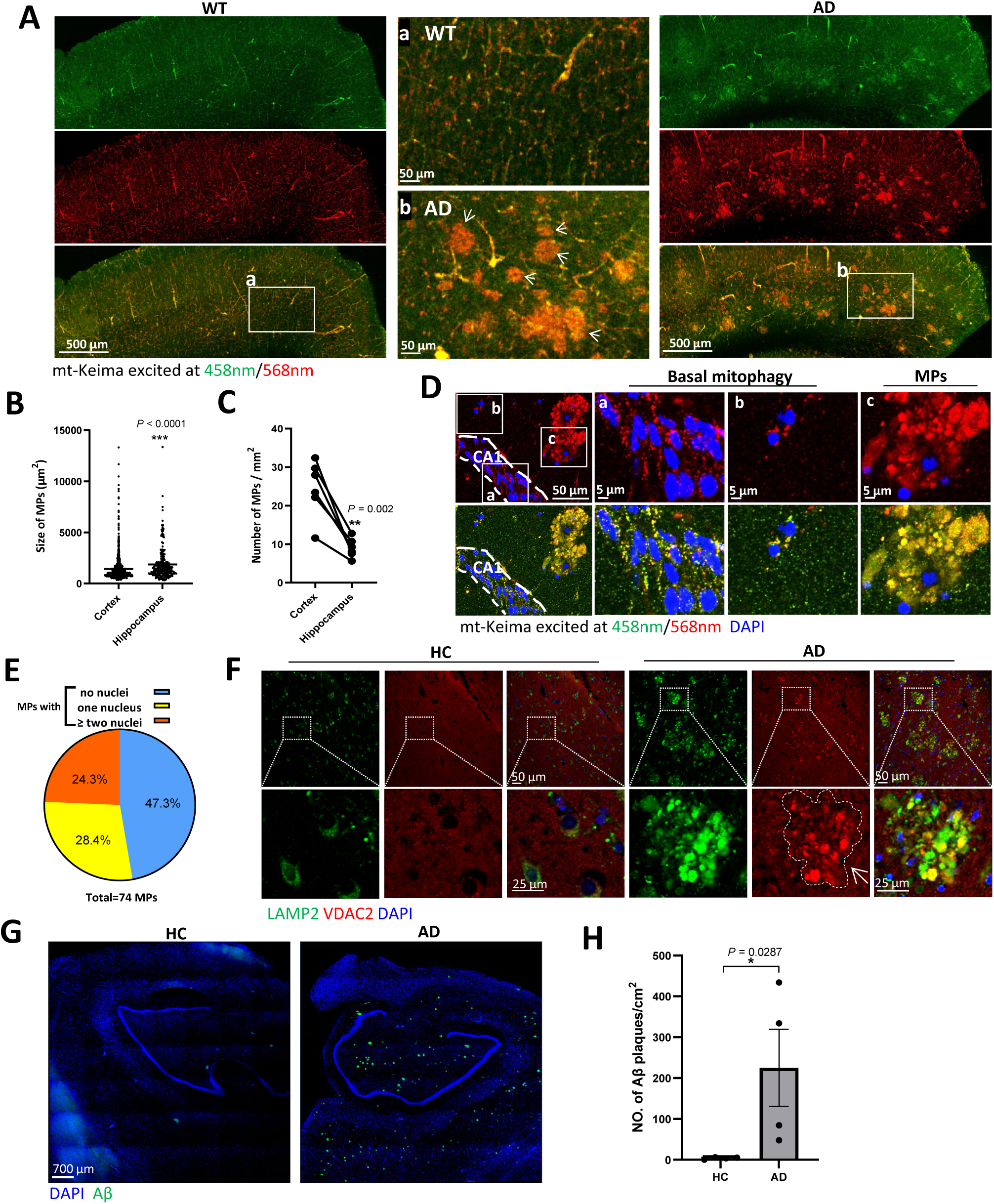
Mitochondrial-enriched structures are specifically detected in AD mice and human brains. (A) Immunofluorescent images showing MPs (white arrows) in cortex of AD mouse brain at 50-60 wks by *ex vivo* imaging. (B-C) MPs sizes in B and density in C in cortex and hippocampus of AD mice; n = 844 MPs in cortex, n = 229 MPs in hippocampus from 6 AD mice at 50-60 wks, including both sexes (two-tailed Mann Whitney test for B; two-tailed paired t-test for C). (D) Comparison of mitophagy signals between MPs and basal mitophagy in the hippocampus region of AD mice at 50-60 wks by *ex vivo* imaging. (E) The pie chart shows the percentage of MPs with different nuclei numbers in cortex and hippocampus of AD mice. n = 74 MPs from 6 AD mice at 50-60 wks; including both sexes. (F) Immunostaining showing the expression of lysosomes (LAMP2) and mitochondria (VDAC2) in the hippocampus from healthy control (HC) and AD patients. (G-H) Aβ immunostaining in HC and AD patient hippocampus. n = 4 individuals/group (unpaired t-test, one tailed). Data are mean and SEM. *P < 0.05, P** < 0.01, ***P < 0.001.

MP-like structures (indicated by the arrow) were also specifically observed in postmortem hippocampal tissue from AD patients, but not in age-matched healthy controls (Fig. 1F, S1C). In healthy controls, mitochondria were typically homogeneously distributed around the nuclei; however, in human AD brains, mitochondria aggregated in a similar form as MPs observed in AD mice. These aggregates largely co-localized with lysosome markers (Fig. 1F), supporting their acidic nature as observed in AD mouse brains. The pathology of AD human brains was confirmed by Aβ staining (Fig. 1G, H).

### Mitochondrial plaques are spatially associated with Aβ plaques

To understand the relationship between MPs and Aβ plaques, we analyzed their spatial distribution utilizing reconstructed 3D images (Fig. 2A) derived from mt-Keima fluorescence and Aβ immunostaining (Fig. S1D). While MPs and Aβ plaques exhibited spatial proximity, they remained distinct entities with an average overlapping ratio of 7.3% (Fig. 2A, B and Video S1). The identification of MPs for 3D analysis is depicted in Fig. S1E. The Aβ plaques interacting with MPs involve two main types: primitive and cored (Fig. 2A), as confirmed by transmission electron microscopy (TEM) analysis showing distinct structures populated with mitochondria (red lined) located around primitive or cored Aβ plaques (yellow lined, Fig. 2C). This spatial relationship was also observed in AD patient brains (Fig. 2D), where mitochondria-enriched structures, indicated by higher fluorescence intensity (Fig. 2E), were found around Aβ plaques.

**Fig. 2.**
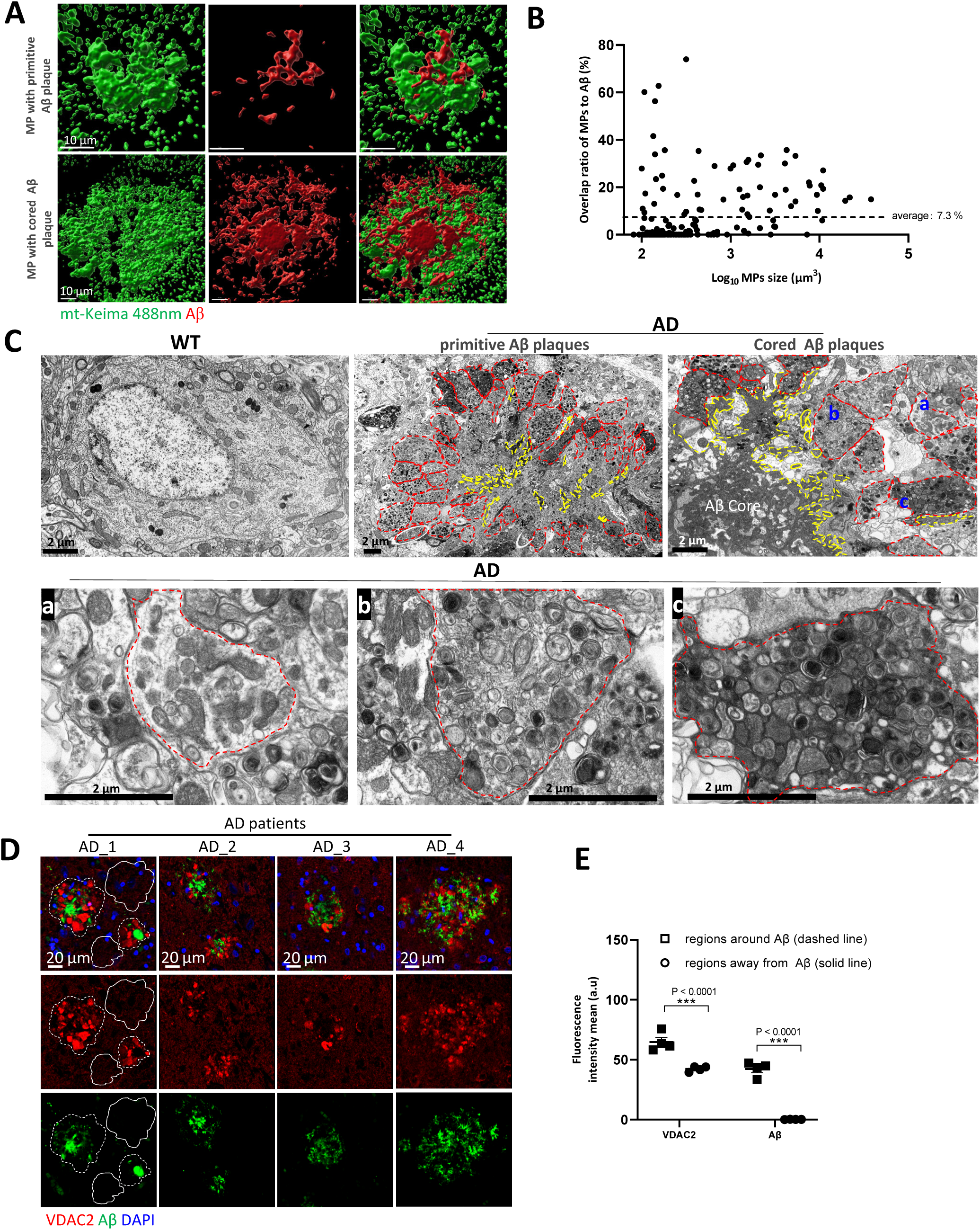
MPs are spatially associated with Aβ plaques. (A) Representative 3D images showing the spatial distribution of MPs and Aꞵ plaques in cortex of AD mice at 50-60 wks. (B) The ratio of overlapping volume of MPs to Aꞵ plaques in cortex of AD mice. n = 196 MPs from 3 AD mice at 50-60 wks, including both sexes. (C) TEM images in the cortex of AD and WT mice at 36-48 wks. Aꞵ plaques and dystrophic neurites are indicated by yellow and red dashed lines, respectively. n = 3 mice/group, including both sexes. (D) Immunostaining on VDAC2 and Aꞵ on the hippocampus from AD patient brains. (E) Comparison of VDAC2 around (dashed lines in D) and away from (solid lines in D) Aꞵ plaques regions in AD patients. n = 4 individuals/group (Two-way ANOVA, Šídák’s multiple comparisons test). Data are mean and SEM. ***P < 0.001.

### MPs harbor amyloid precursor protein and develop into mixed plaques with Aβ plaques

MPs appeared as early as 15 weeks in the cortex with a later onset in the hippocampus between 15 and 25 weeks in our AD mice model (Fig. 3A and S2A). They increased significantly in size and density with age in the cortex region while such changes were less significant in the hippocampal region (Fig. 3A and B). Possibly due to a later onset in hippocampus, a lower MP density was detected than that in the cortex regions (Fig. 3A). Aβ plaques appeared concurrently with MPs at 15 weeks in cortex and increased with AD progression (Fig. 3C). While many MPs contained Aβ (yellow arrows), some MPs lacked Aβ (white arrows) and vice versa (Aβ without MPs indicated by blue arrows; Fig. 3C). Structures with both MPs and Aβ, identified as mixed plaques, consistently increased with age (Fig. 3D, E). MP size positively correlated with Aβ presence and larger MPs are more frequently associated with Aβ (Fig. 3F). This suggests that as MPs grow, they are more likely to develop into mixed plaques with Aβ. The formation of mixed plaques may be attributed to the significant presence of amyloid precursor protein (APP) within MPs. Even though only part of an MP (25%) harbor APP, it accounts for around 60% of APP detected nearby (Fig. 3G, H).

**Fig. 3.**
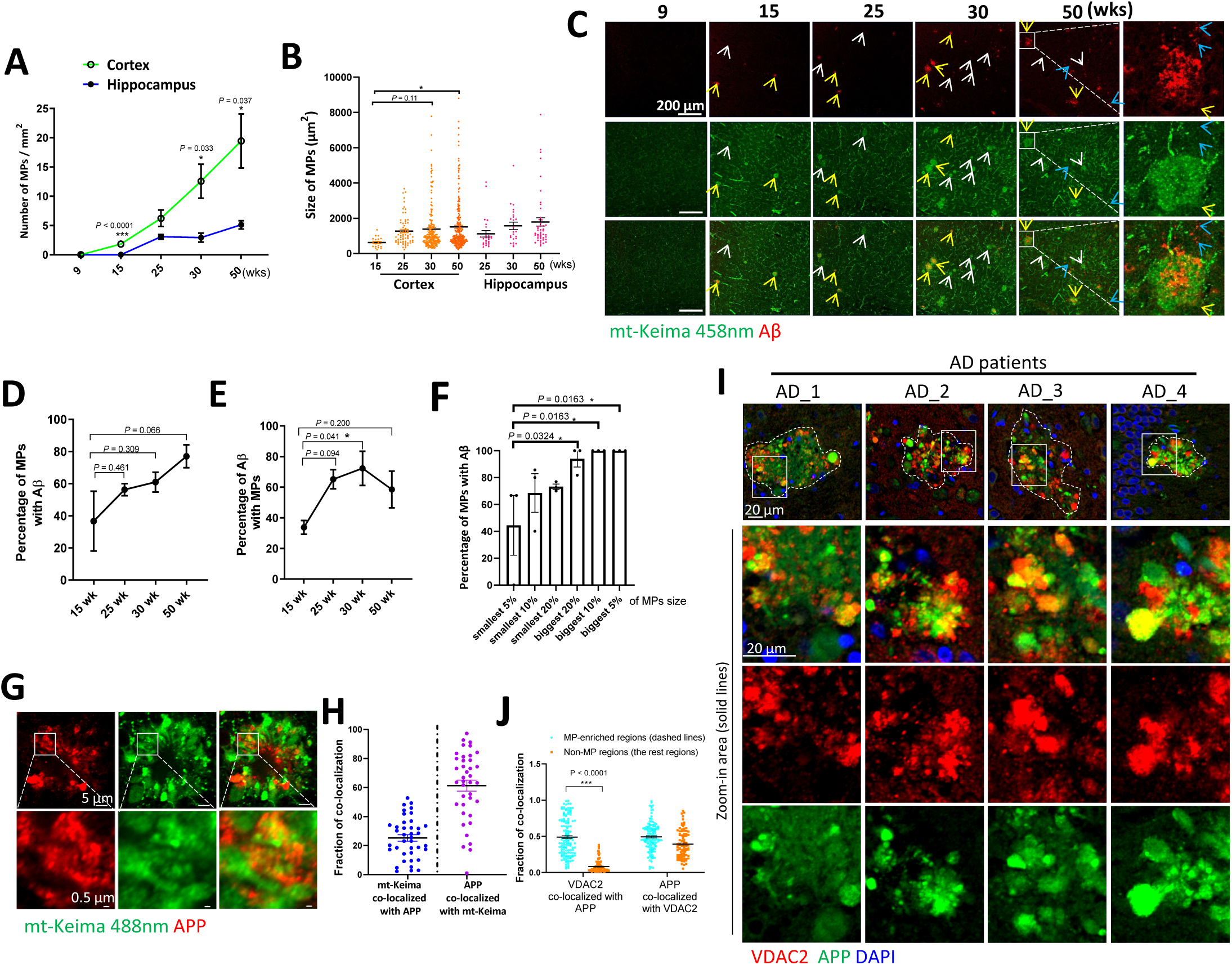
MPs harbor the precursor of Aβ and developed into mixed plaques with Aβ plaques. (A-B) MPs density in A and size in B at different ages of AD mice. n = 3 mice/group, including both sexes (two-tailed unpaired t-tests to compare between age-matched cortex and hippocampus in A; Ordinary one-way ANOVA, Šídák’s multiple comparisons test in B). (C) Fluorescent signals of MPs and Aβ plaques in the cortex of AD mice at different ages. White arrows: MPs without Aβ; yellow arrows: Aβ without MPs; blue arrows: scattered Aβ. (D-E) The percentage of MPs with Aβ in D and percentage of Aβ plaques with MPs in E in the cortex of AD mice at different ages. n = 3 mice/group, including both sexes (One-way ANOVA, Dunnett’s multiple comparisons test). (F) Percentages of MPs with Aβ plaques quantified based on the MPs size percentile in AD mice cortex (50 wks). n = 3 mice/group, including both sexes (one-way ANOVA, Dunnett’s multiple comparisons test). (G) Fluorescent images showing APP and mt-Keima excited at 488nm in a MP region of AD mice at 25 wks (cortex). (H) The co-localization fraction between mt-Keima and APP in MPs enriched regions of AD mice. n = 39 MPs from 3 AD mice at 25 wks, including both sexes. (I) Fluorescent images showing APP and VDAC2 in the hippocampus of AD patients. Dashed lines: MP-enriched regions; solid lines: zoom-in area. (J) Comparison of VDAC2 and APP co-localization between MP-enriched regions and non-MP regions in AD patients. n = 141 and 89 for MPs enriched regions and non-MPs regions from 4 AD individuals, respectively (Two-way ANOVA, Šídák’s multiple comparisons test). Data are mean and SEM. *P < 0.05, P** < 0.01, ***P < 0.001.

Analysis of postmortem human AD hippocampal tissue yielded similar results: around 50% of an MP content harbored APP, while MPs-localized APP constituted 50% of total APP detected nearby (Fig. 3 I and J). Notably, less than 5% of mitochondria in non-MPs regions of AD patient brains were co-localized with APP (Fig. 3J).

### Mitochondrial accumulation in neurites contributes to MPs formation

MPs are distinctly large (Fig. 1B) and contain smaller components (Fig. 1D, 2C, 4A-b) exceeding the size of individual mitochondria (Fig. 4A-a and Video S2) from WT mice (95% CI: 0.166 – 0.172 µm²) and AD mice (95% CI: 0.128 – 0.172 µm², Fig. S2B). We hypothesize that these smaller structures contain accumulated mitochondria, collectively forming MPs.

**Fig. 4.**
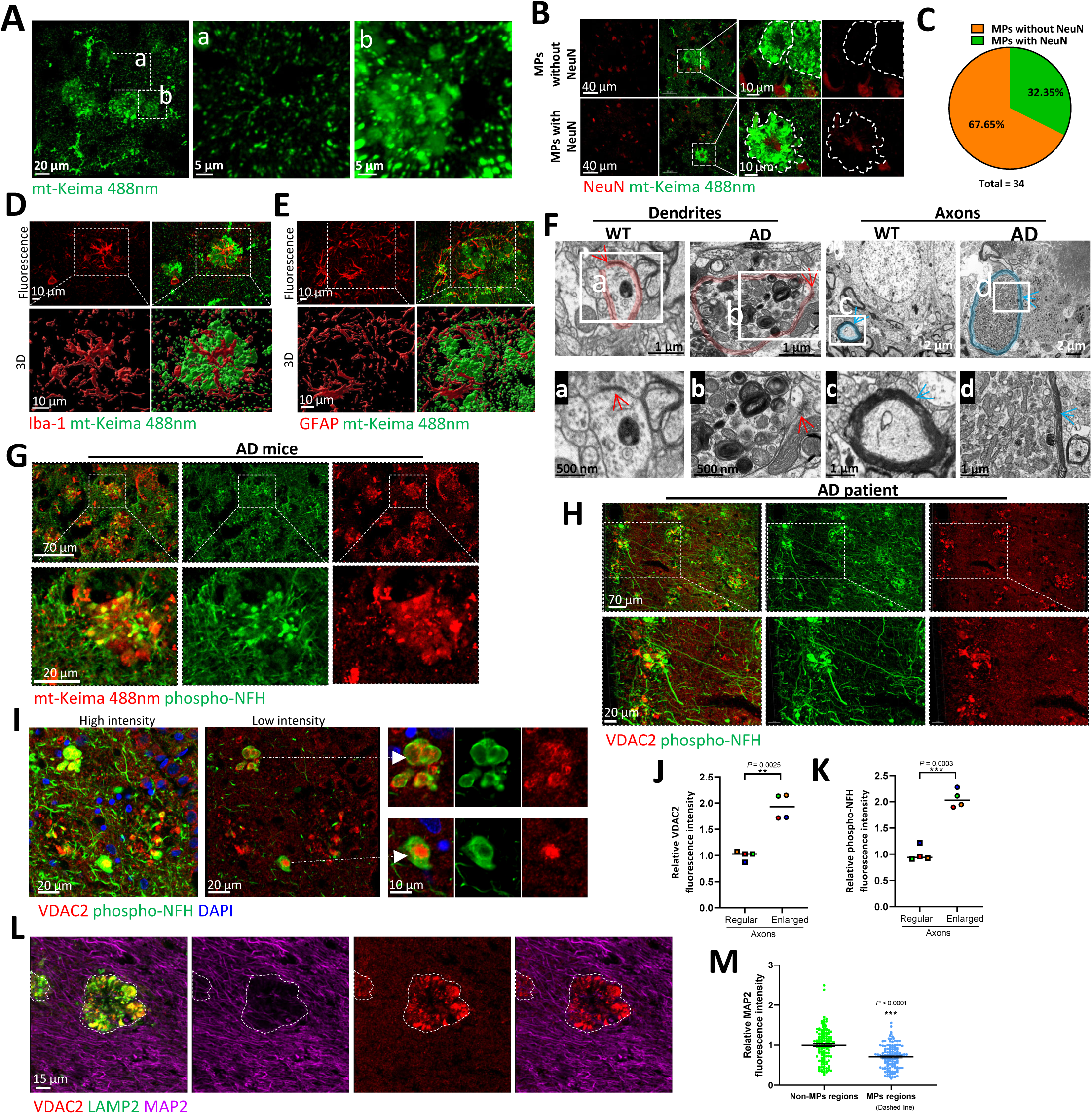
Mitochondria accumulate in dystrophic neurites of AD brains. (A) 3D fluorescent images showing mitochondria (a) and MPs (b) in the cortex of AD mice at 50-60 wks. (B) immunofluorescence images showing MPs (dashed lines) with/without neuronal nucleus in cortex of 25 wks AD mice. (C) The pie chart showing the percentage of MPs with/without NeuN signal. n = 34 MPs from 3 AD mice at 25 wks. (D-E) Immunofluorescent (upper) and re-constructed 3D (lower) images showing the interaction between MPs and microglia in D or GFAP-positive astrocytes in E in cortex of AD at 50-60 wks. (F) TEM images showing the morphology of dendrites and myelinated axons in the cortex of WT and AD mice at 36-48 wks. Red arrows: dendritic spines; blue arrows: myelin sheaths. (G) Immunofluorescent images showing phospho-NFH and mitochondria (mt-Keima 488nm) in the cortex of AD mice at 50-60 wks. (H) Immunofluorescent images showing phospho-NFH and VDAC2 in the hippocampus of AD patients. (I-K) Immunofluorescent images in I and intensity quantifications of VDAC2(J) and phospho-NFH (K) in regular and enlarged axons in the hippocampus of AD patients. n = 4 individuals (paired t-test). (L-M) Immunofluorescent images in L and quantifications in M showing MAP2 expression in MPs regions (dashed lines) and surrounding non-MPs regions in AD patients. n = 132 MPs from 4 AD individuals (two-tailed paired t-test). Data are mean and SEM. P** < 0.01, ***P < 0.001.

To determine whether the structures with accumulated mitochondria are lysosomes near the neuronal nuclei, we stained the AD mouse brains with the neuronal nuclear protein (NeuN). Around 70% of MPs lack NeuN (Fig. 4B, C), suggesting that if neuronal perikarya contributes to MP formation, they represent a small fraction. We then investigated whether MPs consist of accumulated mitochondria from glial cells by observing their spatial relationship. As anticipated, more microglia and GFAP-positive astrocytes were detected in AD mouse brains compared to WT (Fig.S2C, S2D). Notably, these cells closely interact with MPs through penetration (Fig. 4D and Video S3) or encasement (Fig. 4E and Video S4) but remain distinct entities.

Based on these results and TEM characteristics (Fig.2C), the structures with accumulated mitochondria are likely dystrophic neurites (DNs). Supporting this, some of these structures display dendritic spine (red arrow) or myelin sheath (blue arrow, Fig. 4F), indicating dendritic or axonal origins, as referred to TEM anatomy guidance (*19, 20*). This result is further confirmed by immunostaining in both AD mice and patient brains. Enlarged axons stained by phosphorylated high-molecular-weight neurofilament protein (phospho-NFH) contain accumulated mitochondria, as indicated either by aggregated mt-Keima (Fig. 4G) in mice or increased intensity of mitochondrial marker in AD human brains (Fig. 4H-J). The enlarged axons usually showed a higher signal intensity of phospho-NFH than the surrounding normal axons (Fig. 4I, K), suggesting a difference in neurofilament structures important for mitochondrial transport (*21*). It remains unknown whether increased phospho-NFH is a reason or a consequence of mitochondrial accumulation.

In contrast, microtubule-associated protein 2 (MAP2), a dendritic marker, was significantly reduced in MP regions as compared to non-MPs regions in AD mouse brains (Fig. S2E-F) and in patients (Fig. 4L-M, S2G). Despite a low MAP2 signal, MAP2-positive structures were occasionally observed within the MP region with partial co-localization with VDAC2 and LAMP2 (blue arrows, Fig. S2H). This supports the TEM observations regarding mitochondrial accumulation on dendrites (Fig 4F). A lower MAP2 signal in MP regions may possibly be due to dendritic degeneration or dendritic rerouting around MPs. Enlarged axons (Fig. S2I) and the absence of MAP2-positive neuronal processes (Fig. S2G) were only observed in AD mice brain, not in the WT control.

### Mitochondrial accumulation in autophagic vacuoles varies with Alzheimer’s progression

*Ex vivo* imaging of fresh AD mice brains showed heterogeneous acidification of MPs as most of their contents displayed an acidic pH (blue arrows, Fig. S3A), with the rest in a non-acidic environment, as indicated by low signal intensity at mt-Keima 568 nm (white arrows, Fig.S3A). Similarly, around 65% of mitochondria within a MP were co-localized with lysosomal-associated membrane protein 2 (LAMP2) in human AD hippocampus, compared to less than 20% in healthy controls (Fig. S3B-C), suggesting that while most mitochondria in MPs reside in lysosomes, some accumulate outside of them.

We classified mitochondrial-enriched dystrophic neurites (mtDNs) into three categories: Type-I DNs primarily contain mitochondria with few autophagic vehicles; Type-II mtDNs contain both mitochondria and autophagosomes (red arrows); Type-III mtDNs are electron-dense, mainly containing autophagic vesicles and malformed mitochondria (Fig. 5A). Upon closer interrogation, autophagosomes in Type-II mtDNs appear to mainly target mitochondrial degradation, as evidenced by the presence of mitochondria inside (Fig. 5A-a, b, e, f). Type-III mtDNs represent a more mature form containing mainly mitophagosomes (Fig. 5A-c, g) and mitolysosomes (Fig. 5A-d, h). We speculate that mitochondrial accumulation triggers lysosomal recruitment for degradation. Temporal comparison of lysosomal presence in MPs from young to old AD mice shows that much fewer mitochondria within MPs are in lysosomes at 15 wks than 25 wks (Fig. 5B, C), supporting our assertion that lysosomal recruitment is a later response to mitochondrial accumulation. However, with AD progression, this co-localization ratio decreased (Fig. 5B, C), likely due to slower mitochondrial degradation relative to accumulation.

**Fig. 5.**
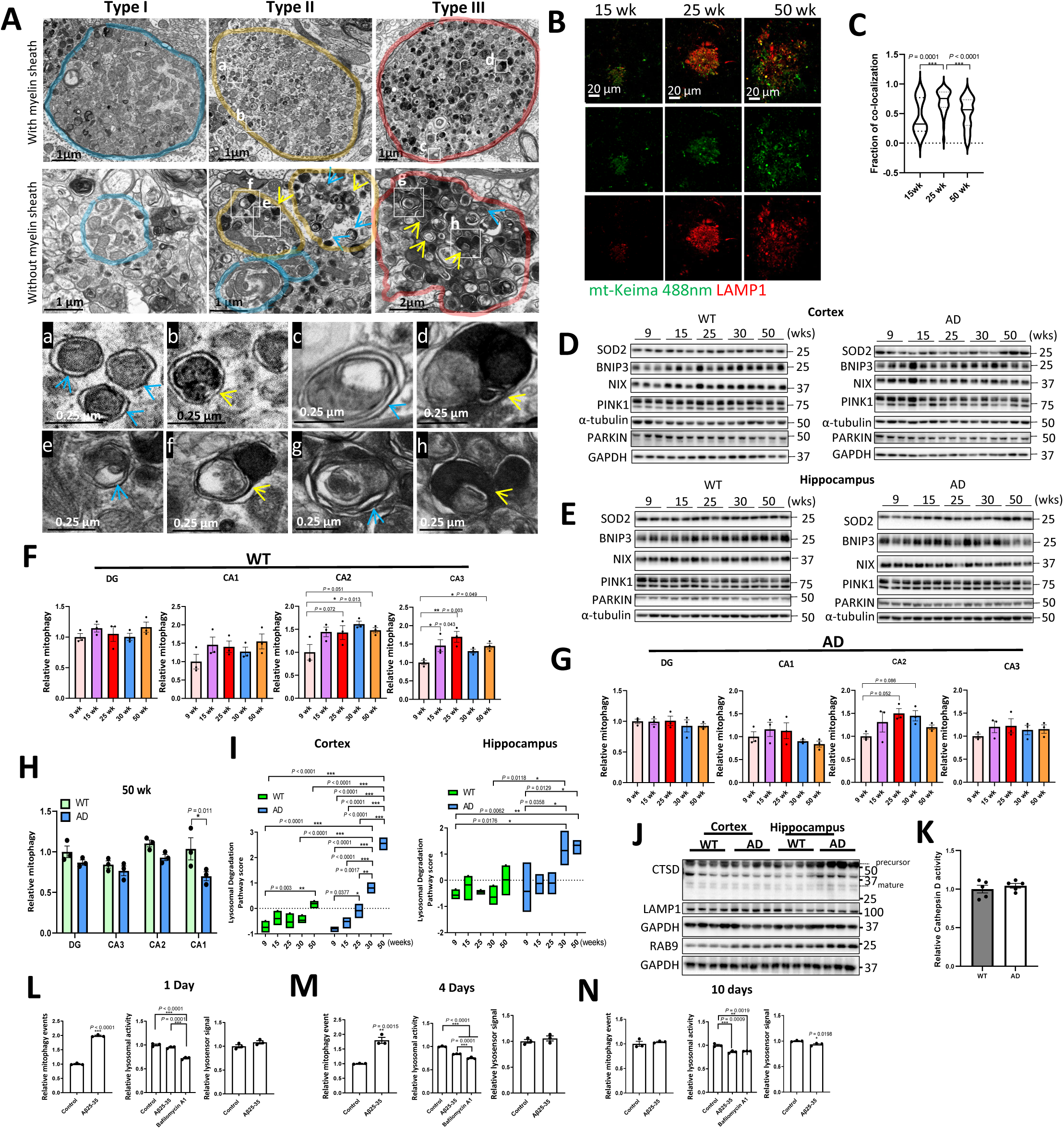
MPs is associated with compromised mitophagy and lysosomal dysfunction. (A) TEM images showing different types of mtDNs in the cortex of AD at 36-48 wks. Type-I, II, and III mtDNs are respectively marked by light blue, orange, and red lines. Blue arrows: mitophagosome; yellow arrows: mitolysosomes. (B-C) immunofluorescence images in B and quantifications in C showing the fraction of a MP co-localized with LAMP1 in the cortex of AD mice. n= 19, 75 and 128 MPs collected from AD mice at 15, 25 and 50 wks, 3 mice/group (Ordinary one-way ANOVA, Tukey’s multiple comparisons test). (D-E) Protein levels of SOD2, BNIP3, NIX, PARKIN and PINK1 in cortex (D) and hippocampus (E) of WT and AD mice from 9 to 50 wks. (F-G) Changes of basal mitophagy across time in different sub-regions of WT (F) and AD (G) hippocampus. n = 3 mice/group (one-way ANOVA, Dunnett’s multiple comparisons test). (H) Comparation of basal mitophagy between AD and WT mice in different hippocampus sub-regions. n = 3 mice/group at 50 wks (two-way ANOVA, Šídák’s multiple comparisons test). (I) Changes of lysosomal degradation pathway score with age in cortex and hippocampus of AD and WT mice. Pathway scores are calculated based on the individual gene expression levels for all the measured genes within a specific pathway for each sample. n = 3 mice/group (two-way ANOVA, Tukey’s multiple comparisons test). (J) Protein expression of CTSD, Rab9 and LAMP1 in WT and AD mice brains (50-60 wks). (K) Relative cathepsin D activity in WT and AD cortex. n = 5 mice/group at 50-60 wks. (L-N) Comparison of mitophagy event, lysosomal activity, and lysosomal pH between SH-SY5Y cells treated without and with 25 µM Aβ25-35 for 24h (L), 4D (M) and 10D (N). n = 3 independent wells. 3 independent biological experiments were done with similar results (two-tailed unpaired t-test or one-way ANOVA, Tukey’s multiple comparisons test). Data are mean and SEM *P < 0.05, **P < 0.01, ***P < 0.001.

### Mitophagy is compromised at the later stage of AD

Given the importance of mitophagy in mitochondrial degradation, we examined how it changes with age in both WT and AD mouse brains. Key regulators for mitophagy, including PINK1, NIX and BNIP3 showed age-related increases in the cortex of WT mice from 9 to 50 wks, a trend absent in AD mice (Fig. 5D, S3D). Notably, BNIP3 increased in the hippocampus of WT mice at 50 weeks, while both NIX and BNIP3 decreased in the AD hippocampus (Fig. 5E, S3E), suggesting a distinct mitophagy regulation pattern between AD and WT mice during aging. Additionally, SOD2, a key regulator of mitochondrial oxidative stress, showed a significant age-related increase in the hippocampus of AD mouse brains, but remained unchanged in WT mice (Fig. 5D-E and S3D-E).

We then directly measured the change of basal mitophagy with age in different sub-regions of hippocampi of AD and WT Keima mice using *ex vivo* imaging. In WT mice, mitophagy increased with age specifically in CA2 and CA3 (Fig. 5F, S3F), but no significant change was observed in AD mice (Fig. 5G, S3F). Notably, CA1 mitophagy was significantly lower in 50-week-old AD mice compared to age matched WT (Fig. 5H), with no significant differences in DG, CA2, and CA3 between age-matched AD and WT mice from 9 to 50 wks (Fig. S3G). These findings, consistent with the results from another older cohort at 50 – 60 wks (Fig. S3H), indicate compromised mitophagy regulation during aging in AD, especially in the CA1 region at the later stage. This may partially explain the decreased lysosomal presence in MPs as AD progresses.

### MP progression is associated with insufficient lysosomal degradation

Mitochondrial degradation is influenced not only by the transport of mitochondria to lysosomes, a process regulated by the mitophagy regulators measured earlier, but also by lysosomal degradation efficiency. Using the nCounter® Mouse Metabolic Pathways Profiling Panel, we found lysosomal degradation to be the most significantly altered pathway in both the cortex and hippocampus of AD mice (Fig. S4A). Notably, these changes were observed at 25 weeks in the cortex and 30 weeks in the hippocampus (Fig. 5I)—both occurring after the initial appearance of MPs in these regions (Fig.3 A), suggesting that lysosomal changes are a response to MP formation.

Neuroinflammatory pathways, including antigen presentation, TLR signaling, NF-κB, and cytokine/chemokine signaling, were altered earlier in the cortex than in the hippocampus (Fig. S4B, C), consistent with the earlier presence of MPs in the cortex of AD mice (Fig. 3A). However, these neuroinflammatory pathways, except TLR signaling in the cortex, showed later onset than lysosomal degradation, suggesting a potential upstream effect of lysosomal dysfunction on neuroinflammation.

Further analysis of lysosomal degradation-related genes revealed increased mRNA expression of key lysosomal proteases (e.g., *Ctsl, Ctsz, Ctsd*, and *Ctss*), lysosomal enzymes (e.g., *Hexa, Hexb, Gusb, Gns*, and *Naglu*), and the glycoprotein CD68 in the AD cortex relative to controls (Fig. S4D), with similar trends but fewer significantly altered genes detected in the hippocampus (Fig. S4E). These results align with the abundant autophagic vacuoles detected in AD brains (*22, 23*).

Recent studies suggest that autophagic vacuoles in AD may be dysfunctional (*11, 13*). To investigate this, we examined the protein expression of Cathepsin D (CTSD), a main lysosomal enzyme, in the cortex and found only the increase of its precursor form, but not the intermediate or mature forms, as compared to the WT control (Fig. 5J, S4F), indicating insufficient maturation. Consistently, no significant differences were observed in CTSD activity between AD and WT mice (Fig. 5K), despite a significant increase in mRNA expression in AD mice (Fig. S4D). Interestingly, all maturation stages of CTSD were significantly higher in the AD hippocampus than WT controls (Fig. 5J, S4F), which may explain the later onset and lower density of MPs in the hippocampus compared to the cortex (Fig. 3A). Additionally, LAMP1 and Rab9 were significantly higher in the AD cortex than in WT controls (Fig. 5J, S4F), confirming the increased formation of autophagic vacuoles in AD. These results suggest that although the number of autophagic vacuoles is increased in response to material accumulation in AD, their actual activity is compromised, resulting in insufficient degradation and accumulation of substrates, like mitochondria.

Supporting this hypothesis, autolysosome acidification was previously shown to be deficient in different AD mouse models (*11*). To further investigate the relationship between Aβ pathology, mitophagy induction, and lysosomal dysfunction, we treated SH-SY5Y cells with Aβ_25-35_. Short-term treatment (1 day) immediately increased mitophagy events (measured using a Dojindo Mtphagy dye) with no significant effects on lysosomal degradation or lysosomal pH (Fig. 5L). Prolonged exposure to Aβ_25-35_ (4 days) led to increased mitophagy events with reduced lysosomal degradation ability and unchanged lysosomal pH (Fig. 5M). Long term treatment (10 days) with Aβ_25-35_ further resulted in defective acidification of lysosomes and dysfunction of lysosomal degradation (Fig. 5N). Notably, the difference in mitophagy events between treated and non-treated cells reduced in the presence of toxic Aβ_25-35_, indicating a negative feedback mechanism of dysfunctional degradation on basal mitophagy following prolonged stress, consistent with our finding of comprised basal mitophagy in the later AD stages (Fig. 5D-G). Altogether, long-term Aβ toxicity results in lysosomal dysfunction, which affects the ability for cargo degradation and contributes to the accumulation of substrates inside.

## Discussion

The present study identified the presence of accumulated mitochondria within or outside lysosomes in both AD mice and human brains, but not in normal controls. We termed these mitochondria-enriched structures mitochondrial plaques (MPs) due to their composition, primarily mitochondria, and their progressive abnormal accumulation like amyloid plaques. MPs can be distinguished from basal mitophagy signals by their characteristic morphology and size. MPs are mainly composed of DNs containing mitochondria (mtDNs) at different stages of degradation, as indicated by TEM in AD mice brains and immunostaining in both AD mice and human brains. Additionally, glial cells were found to infiltrate or encase MPs.

Although MPs may morphologically resemble DNs (*24*) or previously reported lysosome-stained structures (*11*), they are fundamentally distinct from these entities. The previously reported PANTHOS (poisonous anthos (flower)) composed of autophagic vacuoles in AD (*11*) resemble the structures stained by the lysosome markers in the AD mice and human brains in our study. For human AD brains, only around 65% of an MP contents are in lysosomes and only around 60% of a “PANTHOS” structure (stained by lysosome markers) contain mitochondria. This means that while mitochondria are one of the major substrates for “PANTHOS” structure, they are distinct in nature. Notably, approximately 40% of a “PANTHOS” structure in AD human brains lack detectable mitochondria in our study, likely due to severe mitochondrial degradation or presence of non-mitochondrial substrates, like tubular endoplasmic reticulum, axonal cargos, tau, and other proteins, as previously reported (*19, 25, 26*).

MPs are heterogeneous in acidification and content. While MPs mainly consist of mitochondria within lysosomes, some MP contents are simply accumulated mitochondria in neurites (type-I mtDNs). Mitochondrial accumulation may be linked to impaired neuronal cytoskeleton and/or axonal trafficking (*27*). Inhibition of axonal trafficking, including mitochondrial transport, is shown to be an early event in embryonic neurons from PS1 and APP/PS1 mice, with mutation-specific alterations in mitochondrial dynamics preceding AD pathology and amyloid deposition (*28*). Consistently, the levels of kinesin-I and dynein, two microtubule-associated motor proteins were significantly reduced in PS1/APP mouse brains compared to age-matched controls (*14*) and the reduction in kinesin-I is proven to enhance axonal swellings and amyloid deposition (*29*). Additionally, Aβ peptides are also reported to impair mitochondrial transport in neurons (*30*).

The co-localization ratio of MPs with the lysosome marker LAMP1 initially increases with age but later declines in the cortex of AD mice, suggesting an imbalance between mitochondrial accumulation and degradation (mitophagy) as AD progresses. This compromised mitophagy in AD mice is evidenced by age-related changes in key mitophagy-regulating proteins and confirmed through direct *ex vivo* imaging of the fresh brains.

Accumulation of autophagic vacuoles in AD mice brains is observed in our study and in previous publications (*11–13, 19, 29*), however, these autophagic vacuoles may be dysfunctional (*11–14*). Only a few autophagic proteins were detected in DNs of 5xFAD mouse brains while typical autophagosome proteins like P62 and ATG5 were not present (*19*). Cathepsin D, which accumulated in DNs of AD mice, was immature and associated with impaired lysosomal maturation (*31*). In addition, Cathepsin D-positive autophagic vacuoles were poorly acidified in AD mice brains (*11*). In our study, the mRNA expression of Cathepsin D was significantly increased in AD mice cortex, but its activity did not increase accordingly. Defective acidification of lysosomes in response to Aβ toxicity was also observed in our study and this could be a result of direct Aβ toxicity, or a result of progressive accumulation of mitochondrial stress, which overwhelms the degradation machinery. These results together show that while more autophagic vacuoles were formed in AD, their functions are compromised, resulting in the accumulation of autophagic vacuoles containing mitochondria or other substrates. The identification of substrates within these autophagic vacuoles holds value for more specific targeting for AD treatment.

MPs are usually spatially related to Aβ plaques. They start to form at the same time simultaneously or independently at 15 wks, however, with disease progression, they tend to develop into mixed plaques. The precursor form of amyloid is detected in MPs, which may explain such close spatial relationship. Notably, MPs and Aβ plaques remain distinct entities despite their close spatial relations, suggesting the necessity of exploring MPs as a novel and distinct target for AD. Aβ-immunized AD patients revealed successful amyloid plaque clearance yet insufficient protection against ongoing neurodegeneration (*32*). Anti-Aβ plaque interventions can diminish periplaque DNs (*32, 33*), yet persisting DNs (*34*) and mtDNs may still secrete extracellular Aβ. Additionally, MPs mainly affected neurites vital for signal transduction, thus targeting Aβ plaques alone, without tackling these dysfunctional neurites, is unlikely to thoroughly improve cognition decline. Targeting early mitochondrial accumulation and increasing the degradation of accumulated mitochondria would hold promise for future intervention. Our established mouse model would serve as a valuable tool for future intervention studies.

Our studies reveal that mitochondria are one of the primary substrates within autophagic vacuoles in neurites. These lysosome-associated mitochondria, along with other abnormally accumulated mitochondria, contribute to the formation of MPs. Notably, several mitophagy inducers have shown cognitive benefits in AD mouse models, underscoring the critical role of mitophagy (*35*). However, simply enhancing mitochondrial engulfment into autophagosomes may be insufficient, as impaired lysosomal acidification could hinder degradation and exacerbate MP accumulation. Therefore, therapeutic strategies should focus on promoting both mitophagosome formation and lysosomal reacidification. Furthermore, combination treatments targeting both Aβ plaques and MPs to address the mixed plaques hold great promise to provide better therapeutic outcomes in Alzheimer’s disease.

## Acknowledgements and grant support

This work was supported in part by the Intramural Program of the National Institute on Aging, National Institutes of Health (VAB) and U19 AG056278 and U54 AG079754 (PDR). Grant support from NIA also included grant AG000578 to VAB. We thank Dr. Toren Finkel who previously worked at the National Institutes of Health for providing wild-type mt-Keima mice. The APP/PSEN1/mt-keima were established at NIA. We thank Fred Indig of the Confocal Imaging Facility of National Institute on Aging and Patrick Wille, Mary Brown, and Guillermo Marques at University Imaging Centers of the University of Minnesota for providing technique support for confocal imaging. We thank Alfred May at the National Institutes of Health and Ellen Miller at the University of Minnesota for providing technical support. We thank Drs. Simonetta Camandola, Stefano Donega, and Wenlong Liu at the National Institutes of Health for critical reading of the manuscript.

## Data availability

The GSE number for the nano string data is GSE245929. Reasonable requests for all data presented in this article will be honored by the corresponding authors.

## Author contributions

X.D., D.L.C., and V.A.B. designed the study. V.A.B. and P.D.R supervised the project. D.L.C and V.A.B. established the animal model. X.D performed *ex vivo* imaging and X.C. collected the mice samples. X.D. performed the immunostaining and experiments. X.C performed the analysis of TEM images. W.L and X.D performed the analysis of fluorescent images. D.L.C supervised the analysis of Nanostring data. X.D. and X.C wrote and D.L.C., W.L P.D. R and V.A.B. revised the manuscript. All authors contributed to writing the final manuscript.

## List of Supplementary Materials

### Materials and methods

#### Mouse studies

Wild type mt-Keima mice (FVBNCrl.129(Cg)-Igs2^tm1(CAG-mt-Keima)^) were crossed with APP/PS1 double transgenic mice expressing a chimeric mouse/human amyloid precursor protein (Mo/HuAPP695swe) and a mutant human presenilin 1 (PS1-dE9). Mt-Keima mice were backcrossed 5 times onto the 2xAD background. All mice were maintained on a standard NIH diet (T.2018SX.15 Global 18% Protein Extruded Rodent Diet, sterilizable) with food and water available ad libitum. Animals were maintained on a 12-h light/dark cycle, with all testing performed during the light cycle. The animals were group housed, if possible. All animal experiments were performed using protocols approved by the institutional animal care and use committee of the NIA, 361-OSD-2023. Both male and female adult mice were included in the experiments.

#### Human brain samples

Human postmortem hippocampal tissue from AD patients and healthy controls was obtained from the NIH NeuroBioBank. Each group included two males and two females. The ages of the patients ranged from 55 to 66 years, with an average of 62 for the health controls and 63 for the AD group. All AD patients were diagnosed with early-onset Alzheimer’s disease and cerebral amyloid angiopathy, while no clinical brain pathology was observed for health controls based on the information provided by the NIH NeuroBioBank.

#### *Ex vivo* tissue imaging

The protocol of Sun *et al.* for *ex vivo* tissue imaging using a confocal microscope was employed (Zeiss LSM 880) (*16*). Briefly, mice were euthanized by cervical dislocation then brains were immediately dissected and rinsed with cold PBS. The brain was transferred to a pre-chilled Brain Slicer Matrix and coronally sliced into 1-mm sections using the blades. The brain slices were then stained with 5 μg/ml DAPI in cold PBS for 15 min on ice before being transferred to a confocal dish (VWR) for imaging. Based on the pH-sensitive properties of mt-Keima (*16*), dual-excitation ratio imaging in brain tissues expressing mt-Keima was performed with two sequential excitation lasers (458 and 561 nm) in combination with a 570–695-nm emission range (for both excitation wavelengths).

#### Immunofluorescence staining

Fresh mouse brain tissues were fixed in 4% PFA for 24h at 4°C and equilibrated in 30% sucrose for 48h at 4°C. Tissues were then embedded in OCT and sliced at a thickness of 25 µm. For immunostaining with other proteins, brain slices were permeabilized in PBS with 0.3% Triton X-100 and then blocked in 5% BSA with 0.3% TritonX-100 for 1h at RT. Tissue slices were then incubated with primary antibodies diluted in 2.5% BSA with 0.1% TritonX-100 overnight at 4 °C. Slices were washed with PBS and incubated in secondary antibodies diluted in 2.5% for 1 h at RT.

The human brain tissue arrived in 10% neutral-buffered formalin was rinsed with PBS and equilibrated in 15% and 30% sucrose until sinking to the bottom at 4°C. The tissues were then embedded in OCT and sliced at a thickness of 35 µm. Free-floating staining methods were applied after antigen retrieval using 10 mM sodium citrate solution (pH 8.5), permeabilization and blocking in 5% BSA with 0.3% Triton X-100. The sections were treated with TrueBlack® Plus (Biotum, 23014) to reduce autofluorescence after staining.

The specific primary antibodies used for mouse samples include: GFAP (DAKO, Z033401-2), Amyloid (Cell Signaling, 2454s), LAMP1 (Abcam, ab24170), NeuN (Proteintech, 66836-1-Ig), MAP2 (Cell Signaling, 4542), Iba-1 (FUJIFILM, 019-19741), APP (Proteintech, 10524-1-AP). The specific primary antibodies used for human brain samples include APP (Proteintech, 60342-1-Ig), VDAC2 (Proteintech, 11663-1-AP), LAMP2 (Proteintech, 66301-1-Ig), β-Amyloid (D3D2N) (Cell Signaling, 15126), MAP2 (Thermofisher Scientific, PA1-16751). Phospho-NFH (Biolegend, 801602) was applied to both human and mouse samples. The applied secondary antibodies include: ab175652, ab150171, ab175660, ab150113, ab 175471.

Due to the loss of pH gradient during fixation and staining in a neutral pH PBS-based buffer, we collected the mt-Keima signal excited by 458/488 nm lasers for general mitochondria detection in fixed brain tissue, potentially encompassing both intralysosomal and extralysosomal mitochondria. To avoid the interference of the signal of mt-Keima potentially remaining at 568 nm, we avoided using this channel for any immunostaining. All the co-staining of fixed mt-Keima tissues is detected using AlexaFluor 405 nm. Slices were imaged using Zeiss LSM 880 or Nikon A1Rsi Confocal.

#### Image analysis

MPs in 2D fluorescent images are identified as structures enriched with aggregated mitochondria indicated by mt-Keima signal and with a single object sized over 5 μm^2^ which is much larger than a single mitochondrion in mice brain tissues (< 0.2 µm^2^ on average) (*15*).

3D images were reconstructed by the surfaces built based on the absolute fluorescent signal collected by z-stacked images using Imaris 10.0.1 software. A single round mitochondrial object based on a continuous surface built on mt-Keima signal excited at 488nm (fixed tissue) with a threshold of 100 µm^3^ was analyzed as MPs in 3D images. The ratio of MPs overlapped with Aβ in volume was calculated by using object-object statistics based on the constructed 3D surfaces using Imaris. Notably, the cerebrovascular is highly enriched with mitochondria; thus, their reconstructed 3D structures based on mitochondria would likely show a single object surpassing a threshold size while devoid of typical MPs characteristics. Though MPs may also potentially develop in the vascular, single mitochondrial objects with sizes over 100 µm^3^ but in a vascular shape were not considered as typical MPs in the present study. In the 2D images, only a single mitochondria-enriched object sized over 5 μm^2^ in a round shape is measured as an MP.

To compare mitochondrial aggregation in regions near and distant from Aβ plaques in the brains of AD patients, a region of interest (ROIs) surrounding a single Aβ plaque was selected, while a nearby region of similar size, lacking significant Aβ signal, was chosen as the ROI away from Aβ (as shown in the figures). Within each ROI, the average fluorescence intensity of Aβ or VDAC2 in VDAC2-positive structures was quantified using FIJI-ImageJ.

To compare mitochondrial aggregation in enlarged and regular axons in the brains of AD patients, the phosphorylated NF-H stained and balloon-shaped structures with a diameter larger than 2 μm was defined as enlarged axons while those phosphorylated NF-H-stained tubular shape structures with diameter lower than 1 μm are taken as regular axons based on reported diameter of axons in the hippocampus at 0.1 to 1 μm (*36*). The average fluorescent intensity of VDAC2 or phosphorylated NF-H in the defined ROIs was quantified using FIJI-ImageJ.

To assess co-localization between MPs and APP, LAMP1, or LAMP2 in AD mice and/or human brain samples, the mixed structures formed by MPs and APP or LAMP1 or LAMP2 were selected as ROIs for MP-enriched areas (as shown in the corresponding figures), while the remaining image areas were designated as non-MP regions. Co-localization between MPs and the target proteins was analyzed using the JACoP plugin in FIJI-ImageJ. This allowed calculation of the fraction of mitochondrial markers (either mt-Keima or VDAC2) overlapping with the target proteins, as well as the fraction of the target proteins overlapping with mitochondrial markers within the selected ROIs.

#### Cell culture

Human neuroblastoma cell line SH-SY5Y (ATCC) was cultured in DMEM/F12 supplemented with 10% FBS and 1% penicillin–streptomycin at 37 °C in a humidified incubator with 5% CO_2_ with 95% air. SH-SY5Y cells were treated with 25 µM Amyloid β-Protein Fragment 25-35 (Aβ25-35, Sigma, A4559) or ddH2O for 1, 4 or 10 days. Media containing fresh Aβ25-35 was changed every other day.

#### Flow cytometry

To detect mitophagy events produced in 24h, SH-SY5Y cells receiving different treatments were stained with Mtphagy Dye from Dojindo 24h before measurement on flow cytometry. Briefly, cells were washed with FBS-free medium twice before staining with 100 nmol Mtphagy dye in FBS-free medium in the incubator for 30 min. The Mtphagy dye containing medium was moved and cells were washed with FBS-free medium twice before being replaced with medium containing the appropriate treatments. Lysosomal degradation ability was measured using the Lysosomal Intracellular Activity Assay Kit, ab234622, and by following the manufacturer’s protocol. Notably, the cells in the positive control group were treated with DMEM/F12 medium containing 0.5% FBS and 100 nM Bafilomycin A1 1h before incubation with a self-quenched substrate mix. To detect lysosomal pH changes, SH-SY5Y cells receiving different treatments were trypsinized and incubated with an FBS-free medium containing 1 uM LysoSensor™ Green DND-189 (ThermoFisher Scientific, L7535) at 37 °C with 5% CO2 for 30min. Stained cells were kept on ice prior to measurement using flow cytometry (BD LSR II Flow Cytometer). Data was analyzed by FlowJo™ v10.6.1 and the median number was recorded for statistical analysis.

#### Electron microscopy imaging

Six mice, including both females and males (n = 3 per group) were euthanized by cervical dislocation, and the brains were immediately dissected out and rinsed with cold PBS. The brains were transferred to a pre-chilled Brain Slicer Matrix and coronally sliced into 2-mm sections using blazes and fixed in EM Grade Fixative (2.5% glutaraldehyde in PBS) overnight at 4 °C. Brains were rinsed with PBS and cut into smaller pieces before post-fixation in 1.0% Osmium Tetroxide and wash thoroughly in Ultrapure Water. The samples were stained with 2.0% aqueous uranyl acetate, dehydrated in a series of graded ethanols, and infiltrated and embedded in Spurr’s plastic resin. Numerous blocks from each sample were polymerized for 72 hours at 65 C°. Two embedded blocks from each sample were ultra-thin sectioned using a Leica UC7 Ultramicrotome. 60 to 80 nm ultra-thin sections were collected and mounted onto 200 mesh copper grids. The grids were then post-stained with Reynold’s lead citrate and then intensively examined in a FEI Tecnai Spirit Twin Transmission Electron Microscope, operating at 80 kV. Mitochondrial sizes were analyzed using FIJI-ImageJ software.

#### Western blot

The cortex and hippocampus tissues were obtained from both WT and AD mice at different ages and lysed using radioimmunoprecipitation assay (RIPA) buffer (Cell Signaling Technology, MA, USA), supplemented with protease inhibitors (Bimake) and phosphatase inhibitors (Bimake). The lysates were sonicated at power 4 for 10 seconds on ice using an ultrasonic cell disruptor. Subsequently, the samples were centrifuged at 14,000 g for 30 minutes, and the supernatants were collected. The proteins were then separated on 4-12% Bis-Tris gel (Thermo Fisher Scientific) and transferred onto a PVDF membrane. Following this, the membranes were blocked in 5% milk (Bio-Rad) in TBST (20 mM Tris-HCl pH 7.4, 140 mM NaCl, 0.1% Tween® 20) for 1 hour at room temperature. Next, the membranes were incubated with a primary antibody overnight at 4°C. Antibodies included: CTSD (Proteintech, 21327-1-AP), LAMP1 (Abcam, ab24170), Rab 9 (Proteintech, 11420-1-AP) and GAPDH (Cell Signaling, #2118), PINK1 (Cell Signaling, #6946), PARKIN (Proteintech, 14060-1-AP), BNIP3 (Cell Signaling, #44060S), Nix (Cell Signaling, #12396S), α-tubulin (Proteintech, 66031-1-Ig). Afterward, blots were incubated with an HRP-conjugated secondary antibody (Cell Signaling) for 1 hour at room temperature. The signals were then detected using a PharosFX plus Molecular Imager (Bio-Rad), and the band intensity was determined using FIJI-ImageJ software.

#### Measurement of Cathepsin D activity

The activity of Cathepsin D in the cortex tissue was measured by using a Cathepsin D activity assay kit (Abcam, ab65302) following the manufacturer’s protocol.

#### Nanostring and data analysis

Mouse cortex and hippocampus were collected at 9, 15, 25, 30, or 50 wks and snap frozen in liquid nitrogen. RNA was isolated using a PureLink™ RNA Mini Kit (ThermoFisher Scientific, 12183018A) according to the manufacturer’s guide. The quantity and quality of RNA were analyzed using a Nanodrop, ND-100 spectrophotometer. 200 ng RNA per sample was analyzed on the nCounter platform with nCounter (4.0) Mouse Metabolic Pathways Panel which contains 748 genes related to cellular metabolism (NanoString Technologies). For each batch of samples to be analyzed, the raw data was normalized to the geometric mean of the internal positive controls as well as to housekeeping genes selected by nSolver Analysis software. Directed global significance statistics measure the extent to which a gene set’s genes are up- or down-regulated with the variable by using WT at 9 wks as the control. Red and blue denote gene sets whose genes exhibit extensive over-expression or under-expression with the covariate, respectively. Pathway scores are calculated based on the individual gene expression levels for all the measured genes within a specific pathway for each sample.

#### Quantification of basal mitophagy and acidic fraction of MPS

To prevent interference from MPs, regions containing MPs were excluded from basal mitophagy quantification in the hippocampus. Mitophagy was calculated using Zeiss ZEN 3.0 (blue edition) as previously reported (*16*). Briefly, mt-Keima fluorescence signals obtained from excitation with a 561 nm laser (acidic) and a 458 nm laser (neutral pH) were respectively shown in red and in green and plotted in a scatter diagram based on fluorescence intensity. Mitophagy was calculated by dividing the number of pixels with high red intensity by the total number of mitochondrial pixels which is defined as the total pixel number after subtraction for background pixels. Images with the same imaging and analysis parameters were applied for data acquisition and comparison.

#### Quantification of Aβ with/without MPs and MPs with/without Aβ

The area of interest (ROI) for each MP and Aβ plaque were independently defined based on their fluorescence signal. The average fluorescence intensity of MP and Aβ plaque obtained from all samples at different age groups were collected to get the minimum value to set up the threshold value, respectively. To define whether a certain MP is positive of Aβ signal, the ROI was first defined based on the MP signal (ROI-MP), then within this region, the ROI for Aβ plaques (ROI-MP-Aβ) was re-defined based on the Aβ signal. This is because Aβ plaques are often found within MPs and are much smaller than MPs. If the average fluorescence intensity of Aβ in ROI-MP-Aβ is above the set-up threshold for Aβ plaques, this MP is then defined as positive of Aβ plaque, or negative if otherwise. To define whether a certain Aβ plaque is positive of MPs signal, the ROI was first defined based on the Aβ signal (ROI-Aβ), if the average fluorescence intensity of MPs within this region is above the threshold set up for MPs, this Aβ plaque will be defined as positive of MPs, or negative if otherwise. On the scatter plots, the average fluorescent intensity (arbitrary unit) of MPs and Aβ were respectively shown on the X and Y axes. The threshold for each is shown by the dashed lines.

#### Statistical analysis

GraphPad Prism 6.0 was used for statistical analysis. The data are shown as the mean ± SEM with *P < 0.05; **P < 0.01; ***P < 0.001 considered statistically significant. Statistical significances and sample sizes (n) are reported in the figures or figure legends. The student’s T-test was used for comparisons between the two groups. Group differences were analyzed with one-way or two-way ANOVA for comparison among multiple groups.

### Video legends

**Video S1** Representative 3D images showing the spatial distribution of MPs (aggregated green) and Aꞵ plaques (pseudo red) in the cortex region of AD mice at 50-60 wks old. Mitochondria (green) are identified by mt-Keima signal excited at 488 nm using fixed tissue.

**Video S2** Representative 3D fluorescence images showing cell nucleus (blue), MPs and mitochondria in the cortex region of AD mice at 50-60 weeks old. Mitochondria (green) are detected by mt-Keima signal excited at 488 nm using fixed tissue.

**Video S3** Representative 3D images showing the spatial distribution of MPs (aggregated green) and microglia (Iba1 in pseudo red) in the cortex region of AD mice at 50-60 wks old. Mitochondria (green) are identified by mt-Keima signal excited at 488 nm using fixed tissue.

**Video S4** Representative 3D images showing the spatial distribution of MPs (aggregated green) and GFAP positive astrocytes (pseudo red) in the cortex region of AD mice at 50-60 wks old. Mitochondria (green) are identified by mt-Keima signal excited at 488 nm using fixed tissue.

**Fig. S1.**
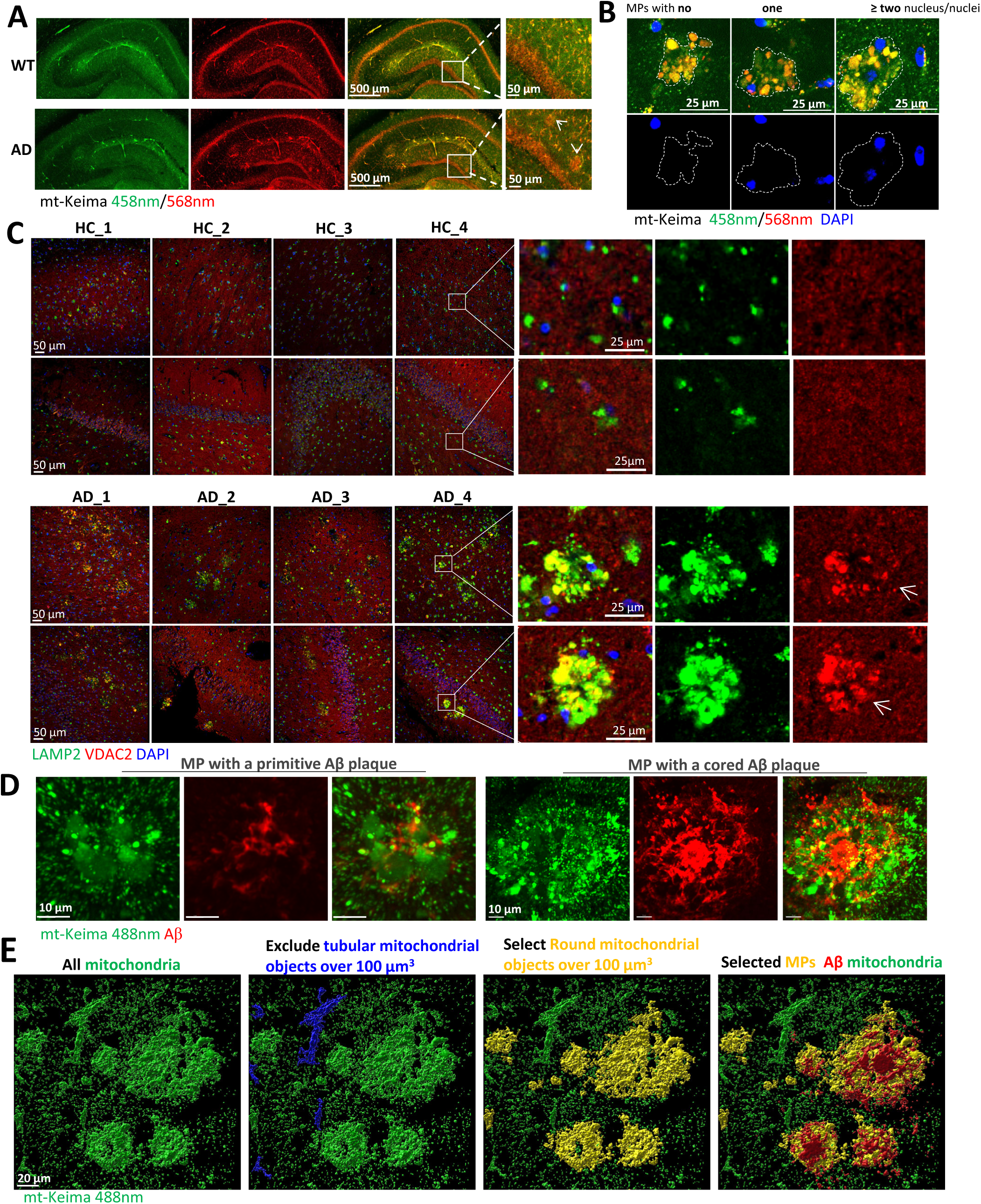
MPs and Aβ plaques develop into mixed plaques. (A) MPs (arrows) in the hippocampus of AD mice at 50-60 wks. (B) MPs (dashed lines) with/without nuclear signal in the hippocampus of AD mice at 50-60 wks. (C) Immunofluorescent staining of lysosomes (LAMP2) and mitochondria (VDAC2) in the hippocampus of HC and AD patients, with MPs indicated by arrows. (D) Fluorescent images showing MPs and Aβ in the cortex of AD mice at 50-60 wks. (E) Reconstructed 3D images depicting the selection of MPs for analysis in cortex region of AD mice at 50-60 wks. Tubular mitochondrial objects over 100 µm^3^ (blue) were excluded from analysis due to the difficulty in distinguishing their classification as either mitochondria-enriched vasculature or MPs (detailed in the method). Round mitochondrial objects over 100 µm^3^ (yellow) are identified as MPs. n = 3 AD mice at 50-60 wks.

**Fig. S2.**
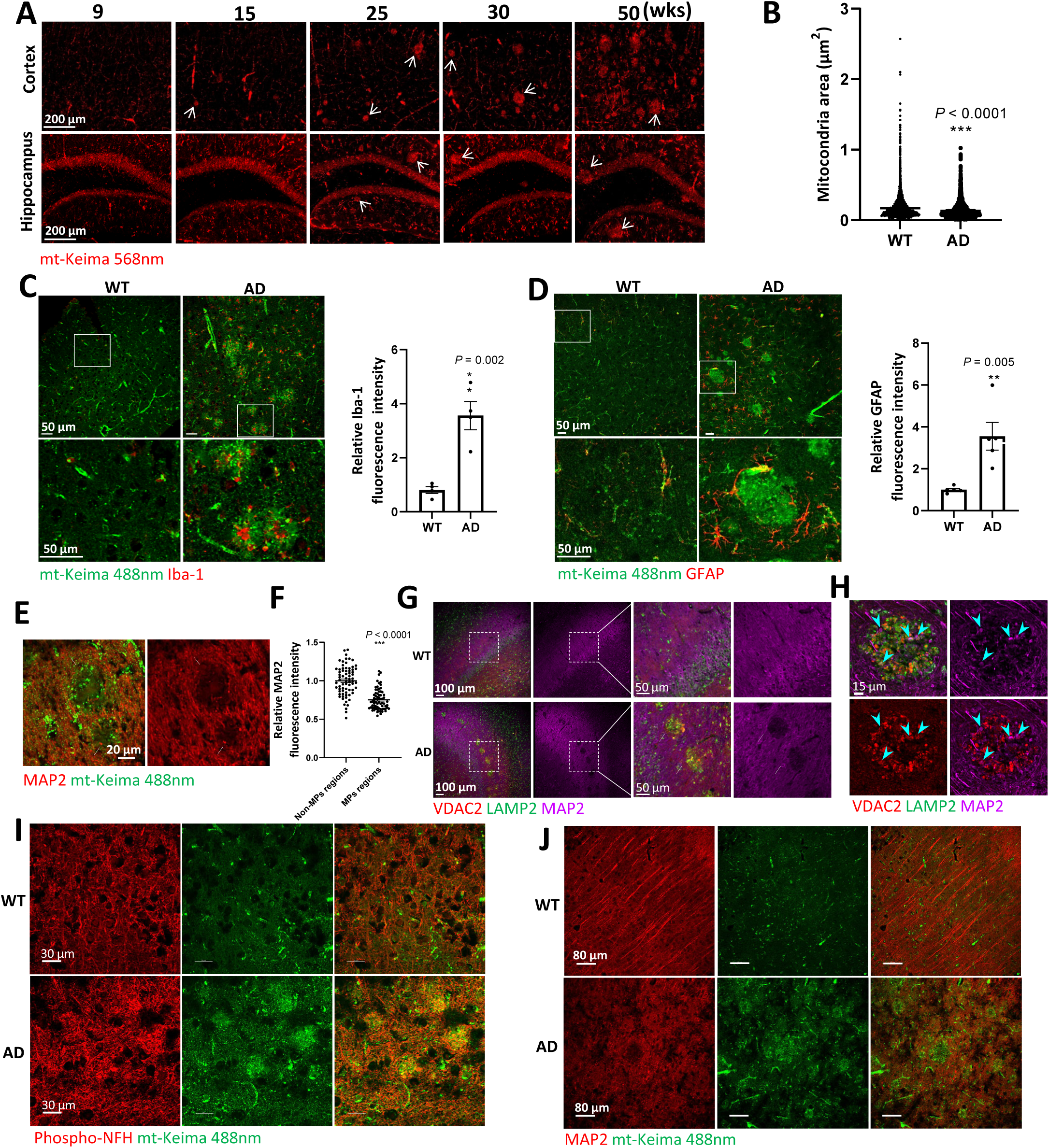
MPs are observed with glial cell recruitment and a lower MAP2 expression. (A) Fluorescent images of MPs (arrows) in cortex and hippocampus of AD mice at different ages. (B) Quantification of mitochondrial area in the cortex of WT and AD mice. n = 8640 and 2926 mitochondria from 3 WT and AD mice at 36-48 wks respectively (two-tailed Mann Whitney test). (C-D) The expression of Iba-1 in C and GFAP in D in the cortex of WT and AD mice at 50-60 wk. n = 4 mice/group in C, n = 5 mice/group in D (unpaired t-test). (E-F) Immunofluorescent images in E showing the MAP2 expression in AD mice brain and comparison of MAP2 signal between the MPs (arrows) and non-MPs regions of AD mice in F. n = 71 for MPs regions or non-MPs regions from 3 AD mice at 50-60 wks (two-tailed unpaired t-test). (G) Immunofluorescent images of MAP2 in the hippocampus of AD and HC individuals, with MPs indicated by arrows. (H) Immunofluorescence images showing the colocalization between MAP2 and VDAC2 in MPs of AD individuals. (I-J) Immunofluorescence images showing phospho-NFH in I and MAP2 in J in the cortex of AD and WT mice at 50-60 wks. Data are mean and SEM. **P < 0.01, ***P < 0.001.

**Fig. S3.**
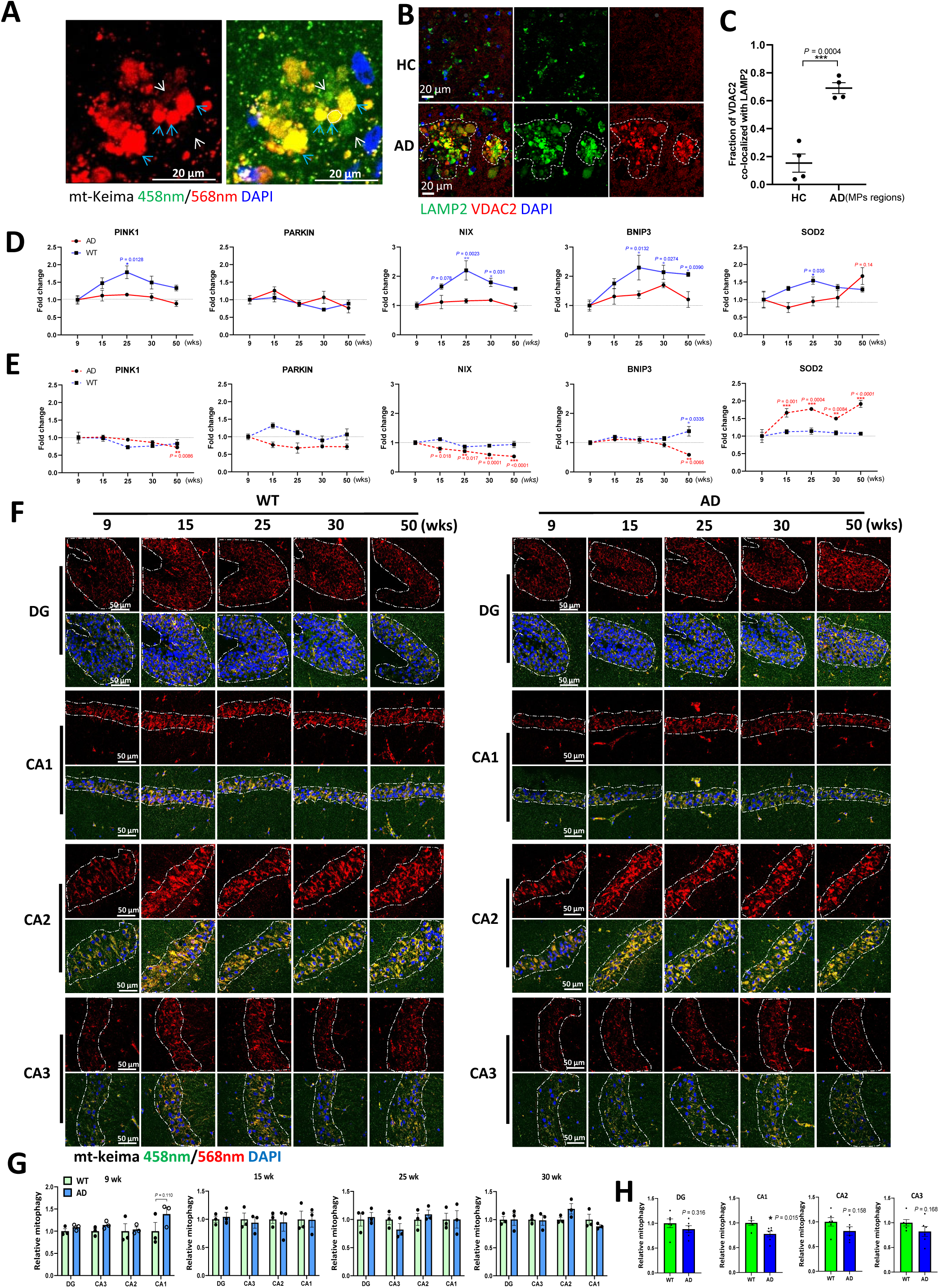
Mitophagy is differentially regulated during aging in AD mice brain. (A) Immunofluorescent images showing acidification of MPs components in the cortex region of AD mice at 50-60 wks. Representative acidic and non-acidic components are indicated by blue and white arrows, respectively. (B-C) Immunofluorescent images in B and quantitation in C showing the co-localization of VDAC2 and LAMP2 in the hippocampus of HC and AD patients, with MPs indicated by dashed lines. n = 4 individuals/group (unpaired t-test). (D-E) Protein levels of PINK1, PARKIN, NIX, BNIP2 and SOD2 in cortex (D) and hippocampus (E) of WT and AD mice from 9 to 50 wks. Statistical comparisons are made by comparing different age groups to the 9-week group for the WT or AD mice respectively, with P value color-coded as indicated. n = 3 mice/group (one-way ANOVA, Dunnett’s multiple comparisons test). (F) Immunofluorescence images showing mitophagy in different sub-regions (white dashed lines) of hippocampus in WT and AD mice at 50-60 wks. (G) Comparation of basal mitophagy between age-matched AD and WT mice in different hippocampus sub-regions from 9-30 wks. n = 3 mice/group (two-way ANOVA, Šídák’s multiple comparisons test). (H) Comparation of basal mitophagy levels between AD and WT mice at 50-60 wks in different sub-regions. n = 6 mice/group (unpaired t-test). Data are mean and SEM. *P < 0.05, P** < 0.01, ***P < 0.001

**Fig. S4.**
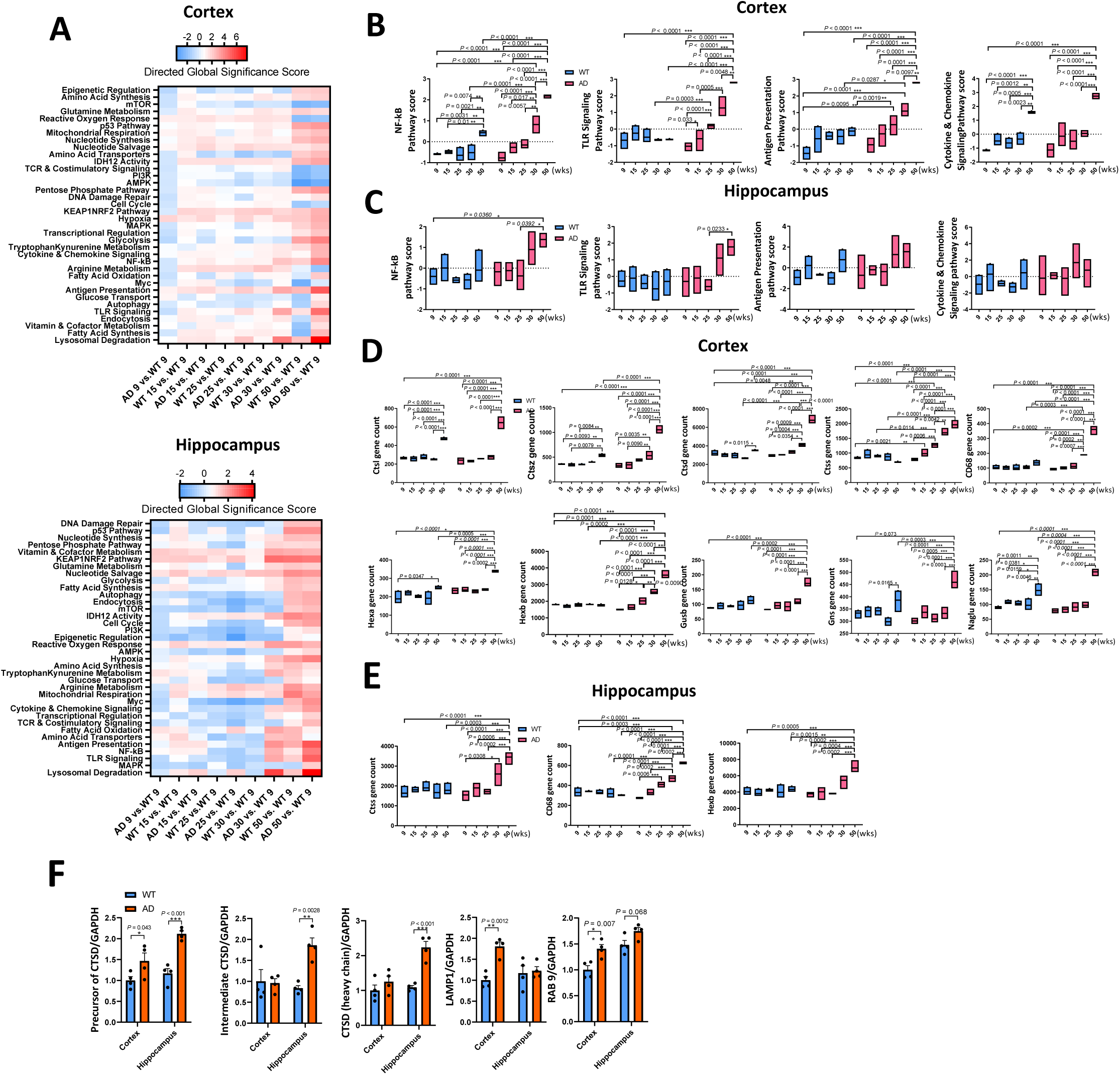
Lysosomal degradation is significantly changed in AD mice brains. (A) The heatmap of global significance scores using nCounter® Mouse Metabolic pathways Profiling Panel on cortex and hippocampus tissue from AD and WT mice aged from 9 to 50 wks. Global significance statistics measure the extent of differential expression of a gene set’s genes with a covariate, ignoring whether each gene within the set is up- or down- regulated. Red and black denote gene sets with extensive and less differential expression with covariance, respectively. n = 3 mice/group. (B-C) Pathway scores on NF-Kb, TLR signaling, and Antigen presentation in cortex (B) and hippocampus (C) of AD and WT mice aged from 9 to 50 wks. n = 3 mice/group (two-way ANOVA, Tukey’s multiple comparisons test). (D-E) Significantly changed genes (normalized gene counts) related to lysosomal degradation in cortex (D) and hippocampus (E) of AD and WT mice aged from 9 to 50 wks. n = 3 mice/group (two-way ANOVA, Tukey’s multiple comparisons test). (F) Quantification of relative protein levels of cathepsin D, LAMP1 and Rab9 using cortex of WT mice at 50-60 wks as reference. n = 4 mice/group (two-way ANOVA, Šídák’s multiple comparisons test). Data are mean and SEM. *P < 0.05, **P < 0.01, ***P < 0.001.

## Notes

### Competing Interest Statement

The authors have declared no competing interest.

